# Ancestral and offspring nutrition interact to affect life history traits in *Drosophila melanogaster*

**DOI:** 10.1101/488809

**Authors:** Joseph B. Deas, Leo Blondel, Cassandra G. Extavour

## Abstract

Ancestral environmental conditions can impact descendant phenotypes through a variety of epigenetic mechanisms. Previous studies on transgenerational effects in *Drosophila melanogaster* suggest that parental nutrition may affect the body size, developmental duration, and egg size of the next generation. However, it is unknown whether these effects on phenotype remain stable across generations, or if specific generations have general responses to ancestral diet. In the current study, we examined the effect on multiple life history phenotypes of changing diet quality across three generations. Our analysis revealed unforeseen patterns in how phenotypes respond to dietary restriction. Our generalized linear model showed that when considering only two generations, offspring phenotypes were primarily affected by their own diet, and to a lesser extent by the diet of their parents or the interaction between the two generations. Surprisingly, however, when considering three generations, offspring phenotypes were primarily impacted by their grandparents’ diet and their own diet. Interactions amongst different generations’ diets affected development time, egg volume, and pupal mass more than ovariole number or wing length. Further, pairwise comparisons of diet groups from the same generation revealed commonalities in strong responses to rich vs. poor diet: ovariole number, pupal mass, and wing length responded more strongly to poor diet than to rich diet, while development time responded strongly to both rich and poor diets. To improve investigations into the mechanisms and consequences of transgenerational, epigenetic inheritance, future studies should closely examine how phenotypes change across a higher number of generations, and consider responses to broader variability in diet treatments.

## INTRODUCTION

For many decades, the consequences of ancestral experiences on the performance and survival of descendants in plants and animals has been a dynamic area of research. Biologists have come to realize that the non-genetic inheritance of environment-dependent effects may represent a significant source of variation for many organismal traits. Recent studies and reviews on humans and mice have highlighted the importance of these phenomena in mediating disease phenotypes, or phenotypes deviating from a defined norm, such as diabetes (Wei et al., 2014), autism (Loke et al., 2015), and cancer (Feinberg et al., 2006). While a substantial portion of effort is directed at piecing together mechanisms of epigenetic gene regulation, biologists have also re-considered the ecological and evolutionary implications of transgenerational effects, by considering the relationship between epigenetic variation and fitness in natural populations (Kilvitis et al., 2014) and how the timing at which a transgenerational effect occurs may determine whether an epigenetic effect is functional versus an impairment (Kuzawa and Thayer, 2011). For example, diversity in the location of methylation marks among populations of bat species may allow them to rapidly buffer change in crowdedness, meteorological conditions (e.g., temperature), noise and light disturbances (Liu et al., 2015). In a second example, early-life grooming of rat pups is associated with changes in methylation of HPA-axis related genes, which are associated with low corticosterone levels and lowered anxiety (Zhang and Meaney, 2010). Female offspring with these modifications groom pups at the same time they receive maternal care, which perpetuates transgenerational transmission of the behavior (Champagne, 2008).

Researchers from various biological disciplines have been compelled to understand the broad and mechanistic significance of ‘epigenetic’ phenomena, and consequently, the literature has been peppered with confusing definitions and co-opted terminology (Haig, 2004). Definitions typically favor a particular organizational level of study, ranging from a strict focus on underlying mechanisms (e.g., DNA methylation, histone acetylation) to the outcome on phenotype (e.g. developmental plasticity; Ho and Burggren, 2010). For the purposes of this study, we use the phrase “transgenerational epigenetic inheritance” to mean any time a form of gene regulation that is not coded by the genomic DNA sequence itself (e.g. factors bound to DNA or freely floating) is inherited by one or more descendant generations, with the inheritance mechanism occurring sometime between germ cell formation and birth of the descendant generation. Different described mechanisms of transgenerational epigenetic inheritance may actually work in concert, so we can imagine a maternal supply of mRNA (influenced or not by the maternal environment), RNA feedback loops, and chromatin modifications as ultimate sources of epigenetic variation that influence phenotypes.

Seven previous studies have, to our knowledge, investigated the patterns of non-genetic inheritance of dietary effects on various traits in *Drosophila melanogaster* (Table 1). Here we briefly review their findings, which suggest some patterns of transgenerational inheritance over one generation, but also leave a number of questions unanswered. Reduction of specific dietary nutrients such as sugar (Buescher et al., 2013; Matzkin et al., 2013), yeast protein (Matzkin et al., 2013; Valtonen et al., 2012) or fat (Dew-Budd et al., 2016), or a dilution of the standard diet (Prasad et al. 2003, Vijendravarma et al. 2010), consistently have effects on egg size, duration of L1 (first instar) -adult development, metabolic pools (concentration of a specific macronutrient in the hemolymph), and body mass in descendant generations. However, the generality of these patterns is questionable, due to differences in experimental design between studies, including wide variability in diet recipes, whether diets differ between parents, and how different diets are between generations. In general, females fed a dilution of standard diet lay larger eggs, from which emerge slowly developing larvae that reach a small body size (Prasad et al., 2003; Vijendravarma et al., 2010). When specific nutrient content is altered, however, these life history traits are differentially affected. For instance, F_1_ females with mothers that ate high protein/low sugar versus low protein/high-sugar were heavier adults, with more protein, glycogen, and triglycerides in their hemolymph, and that laid more eggs, but this pattern changes among genotypes (Matzkin et al., 2013). When sugar content was unchanged in a high versus low fat diet, F_1_ females with mothers that ate high fat food vs low fat food stored fat more rapidly, stored less triacyl glycerides (TAG), had higher concentration of circulating sugars, had increased expression of fat lipolysis and gluconeogenesis genes, and decreased expression of fatty-acid synthesis, sugar transport, and glycolysis genes. These changes in circulating sugar and TAG persisted to the second generation (Buescher et al., 2013). I In the only other study we are aware of in which two generations of potential epigenetic inheritance was considered, the effect of high fat in the grandparental generation on the metabolic pools (macronutrient concentrations in the hemolymph), pupal mass, and egg size of the next two generations, as well as ancestry-independent effects of nutrition, were largely dependent upon genotype and sex (Dew-Budd et al., 2016). In addition, some evidence suggests that offspring ovariole number is influenced by diet restriction in the previous generation: mothers that were deprived of all nutrients birthed daughters that developed more ovarioles than unstarved mothers (Wayne et al., 2006). These studies provided strong support for parental diet influencing some offspring life history traits. However, all but two of these studies (Buescher et al., 2013; Dew-Budd et al., 2016) considered potential effects across only one generation, and none considered the relative strength of a specific generation’s diet versus ancestral diets on different traits.

**Table 1.** Comparison of the reported findings of published articles testing for transgenerational effects of nutrition in life history traits of *Drosophila melanogaster*. The specific composition of “standard diet” differs among studies, and in some cases, is not specified. Ingredients sometimes included in Drosophila media as preservatives, such as MgSO_4_, CaCl_2_, propionic acid, or Nipagin, have also not been included to save space. DGRP: Drosophila Genetic Resource Panel (Mackay et al., 2012). DSPR: Drosophila Synthetic Population Resource (King et al., 2012).

| Number of genetic lines | Generations | Trials | Standard food | Diet treatment | Grandparents diet | Outcomes | Reference |
| --- | --- | --- | --- | --- | --- | --- | --- |
| 1 | F<sub>1</sub> | 1 | Agar-banana-barley flour-jaggery-yeast | Standard,<br>Rich (2X yeast)<br>Poor (no yeast, no jaggery, 2X flour, 2X banana) | Same | Female F<sub>0</sub> fed poor food laid heavier eggs than rich-fed F<sub>0</sub> (insignificant), and had offspring that survived better on rich food. No effects on mass. | (Prasad et al., 2003) |
| 2 lines (one control, one selected for starvation resistance) | F<sub>0</sub>, F<sub>1</sub> | 1 | Cornmeal-molasses-yeast | Standard or no food (water-saturated plug) during adult period | Same | F<sub>0</sub> females (from both control line and starvation resistance line), that were starved as adults for 28 hours, birthed F<sub>1</sub> females that had more ovarioles compared to unstarved F<sub>0</sub> females. | (Wayne et al., 2006) |
| 1 | F<sub>0</sub>, F<sub>1</sub> | 2 | Cornmeal-yeast-sucrose-glucose | Standard<br>Poor (75% less sucrose, 80% less glucose, 75% less dry yeast, 75% less cornmeal) | Same | No diet-mediated parental effects on body size or L1-adult development duration.<br>Females raised on poor food laid larger eggs than females raised on rich food.<br>LP-adult pupal duration unaffected by parental effects or diet | (Vijendravarma et al., 2010) |
| 1 | F<sub>0</sub>, F<sub>1</sub> | 1 | Agar-cornmeal-yeast-syrup | Standard food<br>Poor (-87.5% yeast) | Different and same | Female F<sub>0</sub> fed poor diet birthed larger F<sub>1</sub> females that develop slower. Male F<sub>0</sub> fed poor diet sired larger F<sub>1</sub> males. | (Valtonen et al., 2012) |
| 10 wild-collected isofemale lines | F<sub>0</sub>, F<sub>1</sub> | 1 | Agar-cornmeal-yeast-sucrose | (LP) Low protein (+60% sucrose, -60% less yeast)<br>(HP) High yeast (-60% less sucrose, +60% more yeast) | Same | There were genotype-specific effects, but one prominent pattern was reported: F<sub>1</sub> females from HP parents developed more quickly, were heavier, laid more eggs, and had more standing protein and triglycerides. F<sub>1</sub> males developed at a similar rate, no effects on weight, but had higher standing protein when F<sub>0</sub> parents were LP. | (Matzkin et al., 2013) |
| 1 | F<sub>0</sub>, F<sub>1</sub>, F<sub>2</sub> | 1 | Cornmeal-molasses | (LS) Low sugar (0.15 mol/L)<br>(HS) High sugar (1 mol/L) | Different and same | F<sub>1</sub> male and female F<sub>1</sub> offspring whose mothers ate HS food accumulated triglycerides quickly, stored less cholesterol, and had altered regulation of metabolic genes; these patterns were consistent for F<sub>2</sub>. | (Buescher et al., 2013) |
| 4 and 10 randomly selected DGRP lines; 3 randomly selected DSPR lines | F<sub>0</sub>, F<sub>1</sub>, F<sub>2</sub> | 1 | Cornmeal-molasses | Standard<br>High fat (standard + 3% coconut oil by weight) | Different and same | Genotype-specific effects, for at least one of two descendant generations, of fat treatment on trehalose, protein, and triglyceride levels, female and male pupal weight, and egg size. | (Dew-Budd et al., 2016) (DGRP; DSPR) |

In this study, we were interested in the following questions: How does the effect of parental nutrition change between generations? Are responses of phenotype to diet similar in direction and magnitude? How does two generations of ancestral nutrition affect pupal mass, development duration, ovariole number, egg size, and wing length? We tested the following specific hypotheses about the effects of F_0_ nutrition on F_1_ phenotypes, as predicted by previous studies on *Drosophila melanogaster*: (1) When F_0_ females experience dietary restriction, their descendant generations will lay eggs of increased size, from which emerge more slowly developing larvae, with lower pupal mass. Lower pupal mass may indicate lower ovariole number and smaller wing length, but unknown trait linkages that produce trade-offs between traits may lead to alternative changes in certain phenotypes. (2) When we supplement F_0_ standard food with active yeast, which represented the major protein source in the previous studies, we expect a decrease in F_1_ larval development duration, and an increase in pupal mass.

Our results show support for some of these predictions, but in other cases, we observed unexpected trait- and generation-specific epigenetic phenotypes. Moreover, we found support for unanticipated transgenerational effects on ovariole number and wing length.

## MATERIALS AND METHODS Husbandry

Flies (*D. melanogaster*) used in the experiment originate from an Oregon R-C stock (Bloomington Drosophila Stock Center (BDSC) #5) that was maintained in the laboratory at room temperature (~ 23°C) for approximately 5 years before the study was conducted. Flies were maintained at 25 °C and 60% relative humidity (rh) for the duration of the experiment. Stock larvae were reared on ‘standard’ diet: 8500 milliliters of water, 79 g agar (0.9 % w/v), 275 g torula yeast (3.2 % w/v), 520 g cornmeal (6.1 % w/v), 1100 g dextrose (12.9 % w/v), and 23.8 g methyl p-hydroxybenzoate (an antifungal agent) dissolved in 91.8 mL of 95% ethanol. Experimental larvae of each generation were fed ‘rich’, standard, or ‘poor’ diet. Rich diet consisted of solid standard food supplemented with approximately 30 μl of a torula yeast slurry of density 2.86 μg/μl (1 mg of yeast dissolved in 350 μl water), that was pipetted atop the food. Poor diet was made with freshly cooked standard food diluted with boiled 3% agar in a 1:3 ratio (25% concentration of standard diet) without yeast supplementation.

### Experimental Design

Three groups of 25 females and 14 males were placed in egg collection cages with apple juice plates (90 g agar, 100 g sugar, 1 L apple juice, 3 L water, and 6g Nipagin dissolved in 60 mL ethanol). Each cage consisted of a ventilated 60 mm Petri dish bottom ¾ full of apple juice medium, and a dab of yeast paste. First instar larvae were collected over 3 days, replacing the yeast-supplemented apple juice cage each day. Forty larvae at a time were placed in one vial of a set of 8-10 replicate vials for each dietary treatment. All adults, eggs and larvae were maintained at 25 °C and 60% relative humidity (rh) throughout the experiment.

We examined a total of three generations, recording five life history phenotypes (described in “Phenotype pipeline” below) for each generation, using one or more of three different diets at each generation (described in “Husbandry” above; Fig. 1). The design was aimed at examining the different effects of rich and poor diets in each of the three generations. The effects of “standard” diet were not specifically examined; rather, the standard diet was included only for normalization. Throughout the remainder of the article, we refer to the first of these three generations with the label ‘F_0_’, in some cases referring to them as ‘F_0_ parents’ (of F_1_ offspring), and in other cases, referring to them as ‘grandparents’ (of grandoffspring). We refer to the second generation with the ‘F_1_’ label, considering them either as ‘F_1_ offspring’ (of F_0_ parents) or as ‘parents’ (of F_2_ offspring), and the third generation with the ‘F_2_’ label or ‘grandoffspring’ (Fig. 1C).

**Figure 1.**
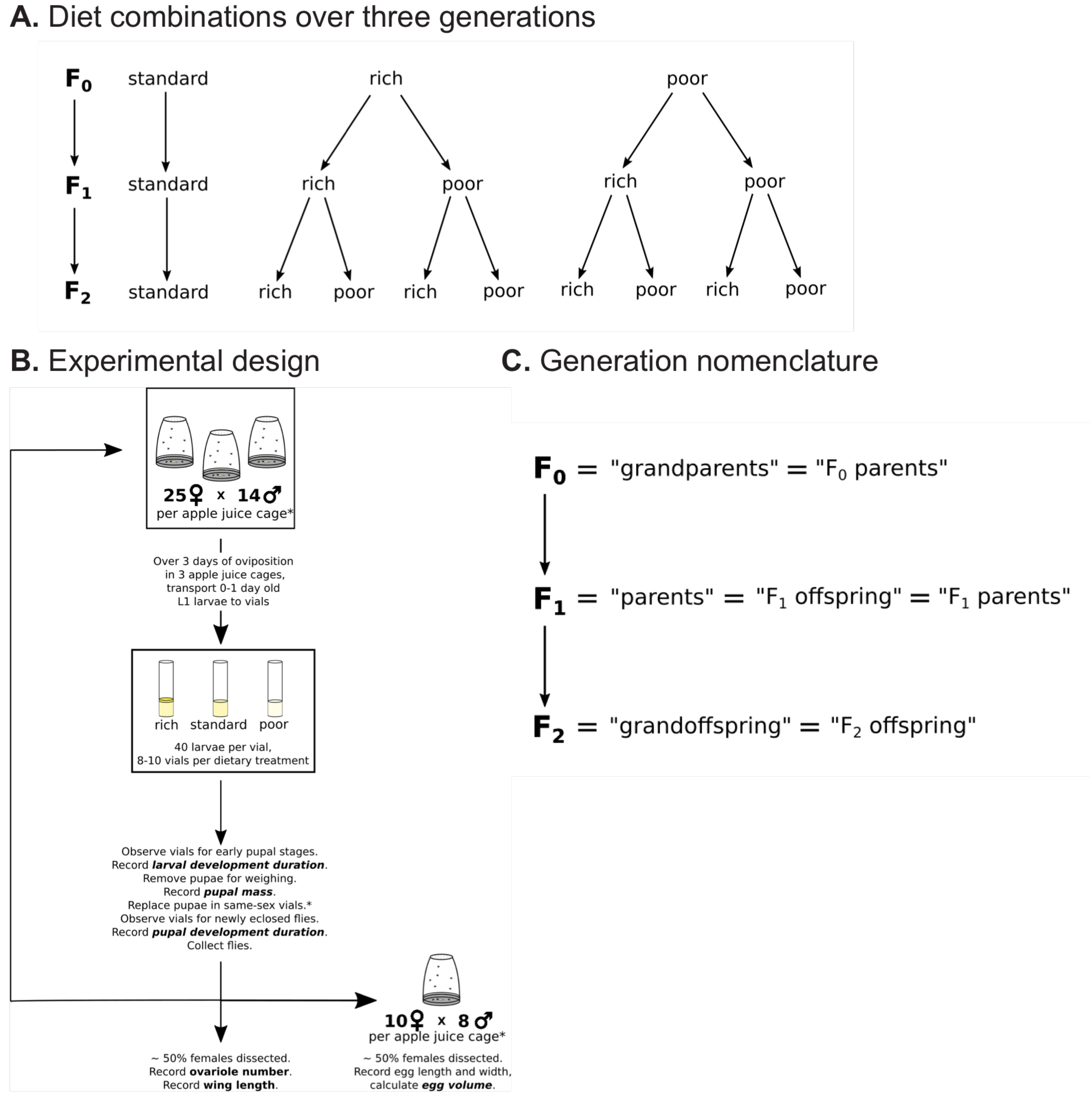
Experimental design and generation nomenclature for the present study. (A) Three generations of flies were observed, subjected to various combinations of diets across each generation. (B) Schematic of the experimental setup for a given generation. See Methods for details. (C) Nomenclature used herein to refer to each generation of the study.

### Phenotype pipeline

The metamorphosis from first instar to adult was recorded every 12 hours. We binned the entire developmental period into three phenotypes, *larval development (L1-LP), pupal development (LP-Adult)*, and *L1-Adult development*. Pupae were sexed, and individual female *pupal mass* recorded, then all pupae were placed into fresh vials and allowed to develop to eclosion.

Upon eclosion, females were kept separated from males in standard food vials supplemented with yeast to stimulate ovariole development and egg production. Three to six days later, approximately 50% of these females were dissected in 1X PBS and their ovaries harvested, which were then fixed in 4% paraformaldehyde in 1X PBS, then stored in methanol at −20°C until full ovariole number counts could be made. Heads and thoraxes were stored in tubes of 70% ethanol for later measurements. After one to three months of storage, ovarioles were gradually rehydrated in a mixture of 1X PBS with 0.02 % Triton-X with DAPI stain at a 1:500 dilution of a 10 mg/ml stock solution, then teased apart using minuten pins. *Ovariole number* for a given treatment group was calculated as the average per ovary.

Facing the ventral side of a dissected thorax, we dissected the right wing, and flattened it in a drop of ethanol on a labeled section of microscope slide. Photos of wings were taken using an eyepiece camera (DinoXcope 7023M) placed in the eyepiece of a Zeiss Stemi DV4 stereo microscope at 25X magnification. Wing photos were viewed in DinoXcope software version 1.16 for Mac OS X. *Wing length* was measured as the distance between the humeral-costal break and the end of vein L3 (see Figure 1 of Gilchrist and Partridge, 1999).

*Egg volume* was estimated by inserting the width and length of an egg into an equation for estimated egg volume: (1/6)πW^2^L (Preston, 1974). These values were averaged for each treatment group. From each treatment group, approximately ten adult females and eight adult males, both aged for four days following eclosion in single-sex vials, were placed in caged, apple-juice plates with a smear of yeast paste to mate and lay eggs. Approximately 20 eggs were collected per diet treatment per generation (treatment groups). Photos of eggs were taken using an eyepiece camera (DinoXcope 7023M) placed in the eyepiece of a Zeiss Stemi DV4 stereo microscope at 25X magnification.

Sample sizes for all phenotypes measured at each generation and diet treatment are included in Supplementary Table 16. Raw data for all phenotypes scored are available at https://extavourlab.github.io/TransgenerationalEffectOfNutrition/.

### Statistical Analysis

All statistical analyses were performed using R version 3.4.3 (Team, 2015). For each generation and phenotype, we constructed a general additive model for location scale and shape (‘gamlss’ package; Rigby and Sasinopoulos, 2005) to analyze the interactive effect of a specific generational group’s experimental diet, and its ancestors’ experimental diet, on different phenotypic responses. We used this model because it allowed us to test each of our response distributions against a larger, general family of model distributions. Moreover, under this model data transformation is unnecessary and one is not restricted to subsets of the exponential family of model distributions. Phenotypes of individuals of the F_2_ generation were potentially affected by three generations of diet (F_0_ diet × F_1_ diet × F_2_ diet), those of the F_1_ generation were potentially affected by two generations of diet (F_0_ diet × F_1_ diet), while those of the F_0_ generation were potentially affected by one generation of diet (F_0_ diet). We used the R package ‘fitdistrplus’ (Delignette-Muller and Dutang, 2015) for fitting model distributions against the observed distribution of a specific phenotype and generation. These distributions are listed in Supplementary Table 8. We selected the one with the lowest Akaike information criterion (AIC) criterion value (Akaike, 1973) for use in our GAMLSS model. This was verified by comparing the AIC values of GAMLSS models (with ‘gamlss’ package) using the distributions with the three lowest AIC values. We also checked whether vial identity should be included as a random effect. In most instances, leaving out vial identity produced a lower AIC value, so we left it out.

We also created an interactive heatmap tool that allows users to visualize the significance of pairwise comparisons of trait values between each of the treatment groups. This heatmap can be used to explore patterns of phenotype differences among any desired set of treatment groups. For example, users can visualize the effect of two generations of poor vs rich diet on grandoffspring ovariole number, of an increase or decrease in food quality on larval development duration, or of any other combination of ancestral diets on the trait of interest. The interactive heatmap of pairwise comparisons, as well as the raw data, are freely available at https://extavourlab.github.io/TransgenerationalEffectOfNutrition/.

For each fly of each generation, we normalized each phenotype response by dividing it by the response of the control (standard food) from the same generation (SS for F_1_, SSS for F_2_). We used this ratio as the response variable for our GAMLSS analyses. We then took the base 10 logarithm of this ratio for use in our graphs (Figs. 2-4). Visualizing the normalized responses relative to each other allowed us to see whether there were general differences in treatment effects. The distribution of values for each normalized phenotype were normal or close to normal using Cullen and Frey plots, so we performed linear regressions. We then assessed for statistical significance using Tukey’s test for multiple comparisons using the ‘multcomp’ package (Hothorn et al., 2008).

**Figure 2.**
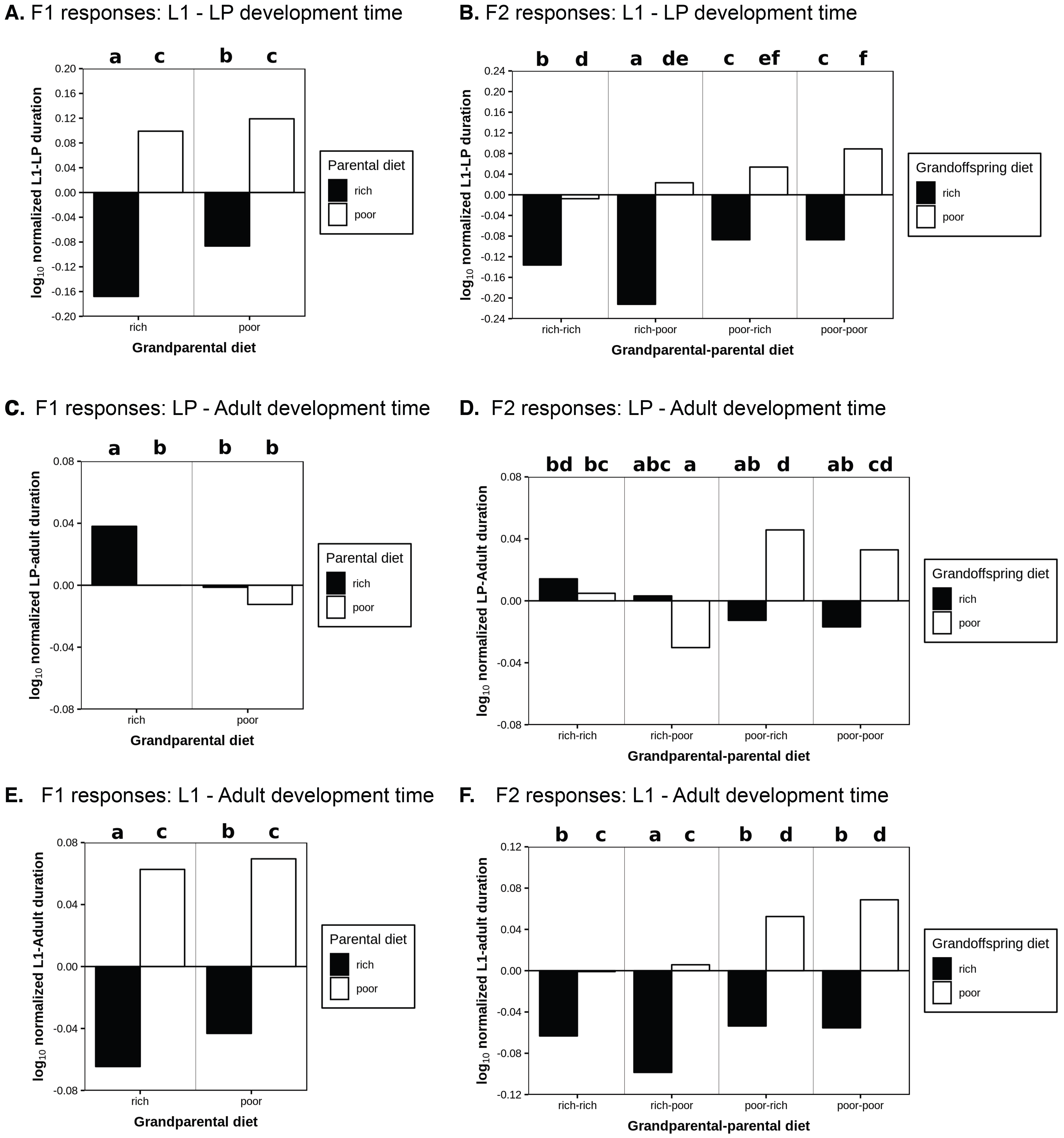
Rich food shortens larval development and lengthens pupal development, regardless of grandparental or grandparental-parental diet. Log-normalized response of developmental duration (y-axis) of parental (A) and grandoffspring (B) generations to larval diet, during the larval (A,B), pupal (A’,B’), and entire developmental period (A’’,B’’). Log-normalization was calculated as the base 10 logarithm of the ratio of parental or grandparental experimental response / control group response (two generations of standard diet for parental, three generations of standard diet for grandparental). Diet treatments are color-coded: rich (black), standard (gray), poor (white). Letters represent how different responses are from each other (using Tukey HSD). Non-significant responses share letters.

## RESULTS

For the three generations examined in this study, for consistency throughout the manuscript we use the naming convention schematized in Figure 1. We refer to the first of these three generations as the F_0_ generation and call them ‘grandparents’ (even when discussing them in relation to their offspring, the F_1_ generation). We refer to the second generation as F_1_, calling them ‘parents’ (of F_2_ offspring). Finally, we refer to the third generation as F_2_ and call them ‘grandoffspring’ (even when discussing them in relation to their parents, the F_1_ generation).

Results of all GAMLSS analyses are reported in Supplementary Tables 1 through 7. We interpreted the regression coefficients from each predictor (F_0_ diet and F_1_ diet for the F_1_ generation; F_0_ diet, F_1_ diet, and F_2_ diet for the F_2_ generation) as the predicted magnitude of change in the value of the measured phenotype. Relative to each other, these coefficients tell us about the relative strength of the effect of the predictor (diet) on a given phenotype. Below, we discuss statistically significant effects revealed by this analysis.

### Parental and grandparental diets have different strengths of effects on offspring phenotypes

We first asked whether ancestral diets, irrespective of their specific nutritional content, showed consistent trends in their influence on distinct phenotypes across two generations. We found that the effects of diet on the parental (F_1_) vs. the grandoffspring (F_2_) generations were distinct from each other and consistent across phenotypes (Suppl. Tables 1-7). Overall, we found that the strongest predictor of most phenotypes for the parental generation was an individual’s own diet, and to a lesser extent, the diet of their parents or the combination of parental and grandparental diets. Contrastingly, we found that for grandoffspring, effects on phenotype were due to their own diet as well as grandparental diet, or interactions among the diets of all three generations. Parental diet alone played little to no role in phenotype effects. Some additional aspects of ancestral diet effects are described in the Supplementary Material (see ‘ Supplementary Results: GAMLSS’).

### Life history phenotypes respond differently to distinct ancestral diet qualities

Next, we asked if life history phenotypes responded to specific combinations of ancestral poor or rich food, relative to conditions where all generations ate standard food. To do this, we normalized the phenotype responses to phenotype values on standard food and log-transformed them, which better revealed differences in how certain phenotypes responded to poor versus rich food. As discussed below, we found that specific hierarchical patterns of diet and ancestry that led to transgenerational effects were not always consistent for rich-or poor-fed parents or grandoffspring, or across generations for the same phenotype.

Overall, this analysis of normalized phenotypes revealed that when compared to poor-fed grandparents, rich-fed grandparents produced parents with shorter larval (Fig. 2A, Suppl. Table 9) and longer pupal development (Fig. 2C, Suppl. Table 10) when those parents were also rich-fed. When poor-fed, parents of rich-fed grandparents had longer wings (Fig. 3A, Suppl. Table 12). Relative to control flies, there was a general trend for F_1_ flies to have lower ovariole numbers (Fig. 4A, Suppl. Table 14) and larger egg volumes regardless of their diet (Fig. 4C, Suppl. Table 15).

**Figure 3.**
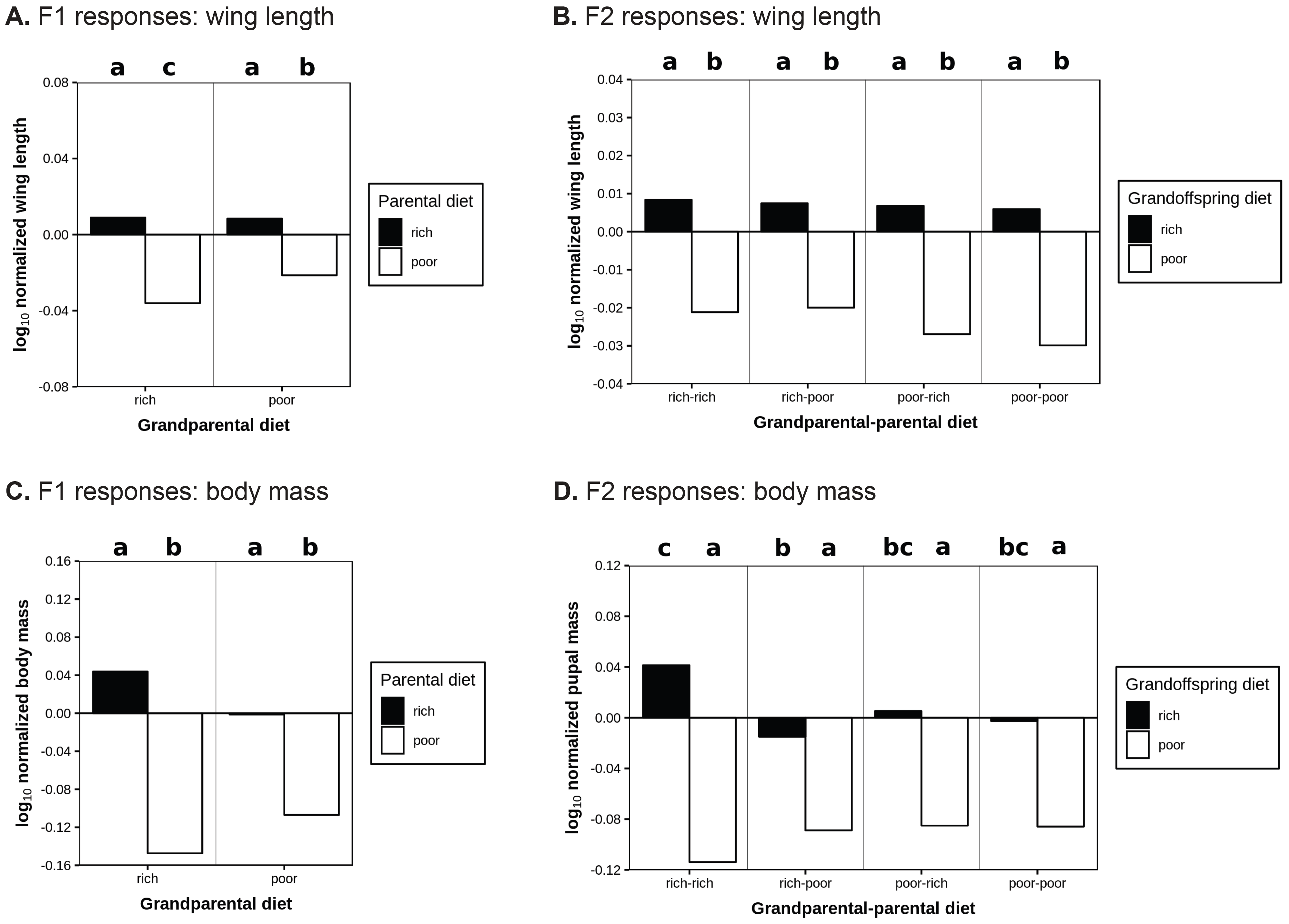
Poor diet more influential than rich diet in changing wing length across two generations; poor-fed flies from otherwise strictly rich-fed lineages have the lowest pupal mass at each generation. Response of wing length of parental (A) and grandoffspring (B) generations to larval diet. Response of pupal mass of parental (C) and grandoffspring (D) generations to larval diet. Log-normalization was calculated as in Figure 1. Color-coding and letters of significance were set as in Figure 1.

**Figure 4.**
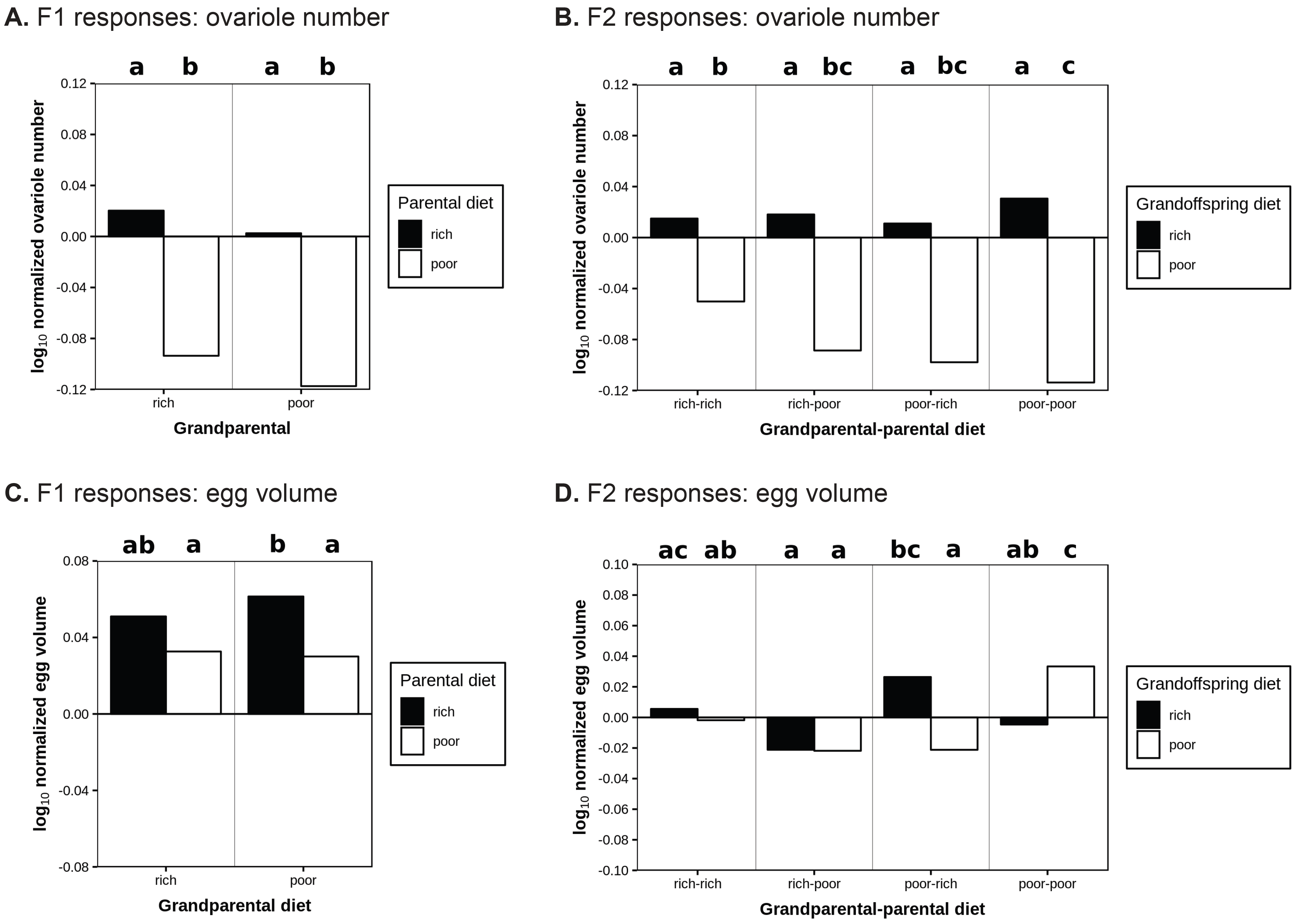
Ovariole number lowered by ancestral poor diets; switching diets among generations causes disordered effects of diet on egg volume. Response of ovariole number of parental (A) and grandoffspring (B) generations to larval diet. Response of egg volume of parental (C) and grandoffspring (D) generations to larval diet. Log-normalization was calculated as in Figure 1. Color-coding and letters of significance were set as in Figure 1.

We found that rich diet in grandparents (F_0_) could often improve fitness-related phenotypes for grandoffspring (F_2_). These phenotypic effects were particularly notable when the parents (F_1_) of those grandoffspring were poor-fed. For example, poor-fed grandoffspring developed more quickly (Fig. 2B, 2D, 2F; Suppl. Table 9-11) when their grandparents were rich-fed than if their grandparents were poor-fed, regardless of the diet of their parents. In other words, a rich diet in grandparents could increase development time of poor-fed grandoffspring, regardless of the parental diet. For rich-fed grandoffspring, if their parents were rich-fed, the grandparental diet did not affect development time one way or another. However, if their parents were poor-fed, then a rich grandparental diet increased larval and overall development time relative to a poor grandparental diet (Fig. 2B, 2F; Suppl. Table 9, 11). With respect to morphological phenotypes, poor-fed grandchildren descended from poor-fed parents had larger wings (Fig. 3B; Suppl. Table 12) and laid smaller eggs (Fig. 4D; Suppl. Table 15) when descended from rich-fed grandparents than from poor-fed grandparents. Two generations of rich-fed ancestors yielded grandoffspring that developed more quickly (Fig. 3B; Suppl. Table 12) and had more ovarioles (Fig. 4B; Suppl. Table 14) than grandoffspring descended from two generations of poor-fed ancestors.

In summary, our study uniquely reveals clear differences in how the phenotypes of each generation are affected by different predictors, and correlative, transgenerational effects of nutrition on sets of phenotypes. Addressing our predictions outlined in the introduction, when grandparents were poor-fed vs. rich-fed, parents did not lay eggs of significantly increased size (Fig. 4C, Suppl. Table 15). Poor-fed vs. rich-fed parents laid significantly smaller eggs, but only if they had poor-fed grandparents (Fig. 4C, Suppl. Table 14). They also had slower development (but only among rich-fed parents; Fig. 2A, 2C, 2E, Suppl. Table 9-11), and shorter wing length (but only among poor-fed parents; 3B, Suppl. Table 12). Ovariole number was statistically invariable among parents (Fig. 4A, Suppl. Table 14). Poor-fed grandoffspring laid larger eggs (Fig. 4D, Suppl. Table 15), had larger pupal mass (Fig. 3D, Suppl. Table 13), lower ovariole number (Fig. 4B, Suppl. Table 14), longer development duration (Fig. 2F, Suppl. Table 11), and shorter wings (Fig. 3B, Suppl. Table 12) if their F_0_ grandparents were poor-fed vs. rich-fed. Rich-fed grandoffspring had longer development duration (Fig. 2F, Suppl. Table 11), and decreased pupal mass if their grandparents were poor-fed vs. rich-fed (Fig. 3D, Suppl. Table 13).

## DISCUSSION

Our GAMLSS results showed that F1 phenotypes were primarily affected by their own diet, and to a lesser extent, the diet of their parents or the interaction between the diets of the two generations. However, F2 phenotypes were primarily impacted by their grandparents’ diet and their own diet. F1 diet individually had little effect on larval and overall development, and no effect on other phenotypes. Interactions amongst the three diet/generation groups had stronger effects on development time, egg volume, and pupal mass than on ovariole number or wing length. When we examined differences among diet groups from the same generation using pairwise comparisons, we found that rich grandparental diet generally decreased the development time, increased the wing length and decreased the egg size of grandoffspring. Two generations of rich diet yielded grandoffspring that developed more quickly and had more ovarioles. This illustrates a difference in the source of phenotypic variation between the parental and grandoffspring generations that is not the same for every phenotype.

We hypothesized that rich vs. poor diets, at least for grandoffspring, interact differently for different traits with the diets of their grandparents. Poor-fed grandoffspring emerged from smaller eggs, laid smaller eggs, had smaller pupal mass, higher ovariole number, shorter development duration, and longer wings if their grandparents were rich-fed vs. poor-fed. Rich-fed grandoffspring had shorter development duration, and increased pupal mass if their grandparents were rich-fed vs. poor-fed. Our study is the first, to our knowledge, to reveal these differences in the effect of ancestral nutrition on multiple life history phenotypes among multiple generations.

We found that transgenerational effects were distinct across generations for different phenotypes. For example, we found that poor-fed grandoffspring had higher ovariole numbers when their grandparents were rich-fed vs. poor-fed. This does not agree with a previous report that food deprivation in previous generations (at least one) increases ovariole number in the next generation (Wayne et al., 2006). A likely explanation for the discrepancy between our results and those of Wayne and colleagues (2006) is the use of different genotypes. This explanation is consistent with the observations of Matzkin and colleagues (2013) and Dew-Budd and colleagues (2016), who both found that the magnitude and direction of effect of ancestral diet varied by sex and genotype. Another possibility is the difference in how we imposed our manipulations. Wayne and colleagues (2006) imposed starvation specifically in the adult generation after larvae had developed on standard food, while we imposed starvation during larval development. This difference in experimental design would be expected to yield different effects on ovariole number, since larval nutrition has been shown to be a major contributor to ovariole number in multiple previous studies (Green II and Extavour, 2014; Hodin and Riddiford, 2000; Mendes and Mirth, 2016; Sarikaya et al., 2012). Wayne and colleagues (2006) also imposed starvation selection on both their “selected” and “control” lines (as opposed to imposing selection on just one line), which may explain why they found effects of starvation on ovariole number in both lines. Lastly, food deprivation in their study consisted of food absence for a specific duration as adults, whereas in our study food deprivation represented a consistent dilution of diet over the entire L1-adult experimental period. Whether food is absent for a short period or in diluted quantities over a long period is likely to interact with the time-specific development of different phenotypes. If development duration is affected in any way by either manipulation, then this may also have cascading effects on other phenotypes.

An under-addressed issue in transgenerational effect studies is how we should interpret dietary manipulations and the effects they have on the organism. Different sugars and yeasts have different effects on life history phenotypes. Flies fed high levels of sucrose suffer reduced fecundity (Bass et al., 2007), while flies fed sucrose but not fructose have reduced lifespan and reduced fecundity (Lushchak et al., 2014), both under normal conditions (Hassett, 1948; Lushchak et al., 2014) and with a starvation period (Hassett, 1948). In *Drosophila* it has been shown that lifespan and fecundity of experimental animals in response to dietary restriction varies depending upon what type of active yeast is used, as well as the amount of carbohydrate relative to protein. This suggests that flies may actually be responding to the restriction of a specific nutrient as opposed to the amount of food (Bass et al., 2007). Likewise, it is possible that the generation-by-diet-dependent changes in egg volume that we observed across generations reflect, in some cases, the rescue of the potential volume (under non-manipulated conditions) from the effects of sub-optimal food types that increase or decrease egg size. To further complicate the issue, there is evidence suggesting that a factor other than amino acids or sugar is present in (at least one species of) yeast, and that this factor is a major nutrient cue for the insulin and TOR signaling pathways that are responsible for nutrient-dependent growth and development (Nagarajan and Grewal, 2014).

One important model for measuring the impact of dietary components on an organism is the geometric framework of nutrition. Nutritional geometry is used to test how combinations of nutrients influence phenotype outcomes, as opposed to considering a specific nutrient in isolation. Principle findings to date suggest that each animal has an intake target (IT), which is the amount and balance of protein and carbohydrates that an animal needs to consume within a specific period to achieve maximal fitness (Raubenheimer et al., 2009). Bondurianksy and colleagues (2016) found that maternal protein influenced offspring pupal mass and head length (a secondary sexual trait) in both sons and daughters, and paternal carbohydrate influenced offspring pupal mass and head length differently between sons and daughters. An alternative hypothesis that could account for these observations, however, is that the intake target shifted in offspring as they aged, which influenced foraging preferences and feeding behavior (Paoli, Donley et al. 2014). In our feeding experiments, it is formally possible that quantified phenotypes were altered not by poor diet *per se*, but by a change in feeding rate in response to “missing” or sub-optimal nutrients in the modified diets.

We have been cautious in our language regarding the interpretation of effects on phenotype. When a non-human animal has increased glucose and triacyl glycerides in their hemolymph, larger body size, and/or lower survival, there is a tendency in the literature to explain these results within the context of metabolic disorders in humans (obesity and diabetes) (Graham and Pick, 2017; Hardy et al., 2018; Musselman and Kuhnlein, 2018; Riddle et al., 2018; Teleman et al., 2012). We would argue that these are medical terms with negative health and socioeconomic connotations that could reflect societal biases in how we perceive changes in human physiology, and making evolutionary arguments about relative fitness gains or costs based on such physiological data may not be appropriate.

For example, an increase in adipose tissue or decrease in lifespan as a result of an increase in sugar or fat content in parental or grandparental diets may, in the end, increase the likelihood of reproducing, regardless of or within specific environmental contexts. The existence of similar regulatory machinery across organisms (e.g. Insulin and insulin-like signaling pathways) does not imply that this machinery achieves the same goals for reproductive fitness across different organisms. Kuo and colleagues (2012) found that *chico* and *Akt* mutants had different cuticular hydrocarbon (CHC) profiles and were less attractive to males than control females. Schultzhaus and colleagues (2018) found that males were less attracted to high-fat-fed females, which had different CHC profiles, than they were to low fat-fed females, in both light and dark conditions. These females also had altered CHC profiles. High fat-fed females are also more fecund (Schultzhaus et al., 2017). However, Lin and colleagues (2018) found that females that ate high yeast diet were heavier, more fecund, less immobile, had shorter lifespans, and had different CHC profiles, but were also more attractive to males than low yeast-fed females, or than high yeast-fed mutant females (hypomorphic for insulin peptides) or oenocyte-specific gene disruption of insulin signaling. These studies illustrate that attractiveness of females, as a measure of reproductive fitness, is represented by CHC profile; these profiles are strongly influenced by high yeast or high fat diet, as well as the disruption of insulin signaling. High yeast and high fat both make female flies heavier, immobile, and more fecund, but the former are more “attractive” in this context, meaning likely to be mated. We acknowledge that transgenerational effects mediated by nutrition may be important with regards to metabolic disorders such as diabetes or obesity in humans. However, we see our results as applying to *Drosophila melanogaster* and other organisms with more similar behavior and physiology, and do not believe it would be wise to speculate beyond these life histories to suggest that our results imply anything predictive or proscriptive about human obesity, fitness, or sugar metabolism.

In conclusion, based on the results of our study and previous studies, we suggest three major potential improvements into future investigations into the transgenerational effects of nutrition on *Drosophila* life history traits. First, due to the variation in transgenerational effects after only one generation, measurements should be conducted for two or more generations. Second, while genotype-and sex-specific effects indicate two important sources of phenotypic variation, general principles are difficult to draw from these effects. A helpful experiment would be to run multiple trials of a well-defined dietary regime on both sexes of multiple genotypes (for example, using the DGRP or a similar collection of population variants (Mackay et al., 2012), in order to discover genotypes with reproducible quantitative phenotypes. These “standard” genotypes would then allow different types of experiments focused at lower organizational levels to be compared across studies. Finally, using a geometric framework of nutrition may give us a robust quantification of the combination of protein, fats, and carbohydrates per genotype that are associated with specific quantitative phenotypes, thereby informing us on how nutritional intake targets, and potentially also correspondent foraging behavior, may play a role in mediating the effects of parental nutrition on offspring phenotypes.

## Supporting information

## ACKNOWLEDGMENTS

We thank the Extavour Lab members for their creative and effective suggestions on experimental design and data analysis. This work was supported by NIH grant R01HD073499 to CGE and funds from Harvard University. JD was partially supported by NIH Supplement 3R01HD073499-03S1.

## SUPPLEMENTARY TABLE CAPTIONS

**Supplementary Tables 1-3. The effect of generation and diet on larval (1), pupal (2), and entire (3) developmental duration.** Summary of GAMLSS used to find broad associations between diet, generation, and phenotype. The coefficients provide the relative size of the effect a model term has on the phenotype. The standard error measures how closely the model estimates the coefficient’s unknown value. The t-statistic is the difference of the estimated coefficient and its true value, divided by the standard error. P-value is the probability of obtaining the t-statistic. One level is missing per generation because it represents the base level against which all other levels are compared. Bolded rows are significant effects.

**Supplementary Table 4. The effect of generation and diet on wing length.** Description of statistical terms and interpretations are as described in Supplementary Tables 1-3.

**Supplementary Table 5. The effect of generation and diet on pupal mass.** Description of statistical terms and interpretations are as described in Supplementary Tables 1-3.

**Supplementary Table 6. The effect of generation and diet on ovariole number.** Description of statistical terms and interpretations are as described in Supplementary Tables 1-3.

**Supplementary Table 7. The effect of generation and diet on egg volume.** Description of statistical terms and interpretations are as described in Supplementary Tables 1-3.

**Supplementary Table 8. List of model distributions tested in GAMLSS analyses.** Comparisons between model and observed distributions was done visually using skewness-kurtosis plots and verified with AIC values, both originating from the R package ‘fitdistrplus’.

**Supplementary Tables 9-11. Pairwise comparisons of larval (9), pupal (10), and entire (11) developmental durations from different generation X diet combinations.** Summary of Tukey contrasts. The coefficient represents the degree of difference between the groups being compared. The standard error measures how closely the model estimates the coefficient’s unknown value. The t-statistic is the difference of the estimated coefficient and its true value, divided by the standard error. P-value is the probability of obtaining the t-statistic. Bolded rows are significant effects.

**Supplementary Table 12. Pairwise comparisons of wing length between different generation X diet combinations.** Description of statistical terms and interpretations are as described in the caption for Supplementary Tables 9-11.

**Supplementary Table 13. Pairwise comparisons of pupal mass between different generation X diet combinations.** Description of statistical terms and interpretations are as described in the caption for Supplementary Tables 9-11.

**Supplementary Table 14. Pairwise comparisons of ovariole number between different generation X diet combinations.** Description of statistical terms and interpretations are as described in the caption for Supplementary Tables 9-11.

**Supplementary Table 15. Pairwise comparisons of egg volume between different generation X diet combinations.** Description of statistical terms and interpretations are as described in the caption for Supplementary Tables 9-11.

**Supplementary Table 16. Descriptive statistics for all phenotypes measured for each diet group at each generation.** Sample size, mean, standard deviation and standard error.

